# A transcriptome and literature guided algorithm for reconstruction of pathways to assess activity of telomere maintenance mechanisms

**DOI:** 10.1101/200535

**Authors:** Lilit Nersisyan, Arsen Arakelyan

## Abstract

Activation of telomere maintenance mechanisms (TMMs) is a crucial factor for indefinite proliferation of cancer cells. The most common TMM is based on the action of telomerase, but in some cancers telomeres are elongated via homologous recombination based alternative mechanism (ALT). Despite their importance, little is known about TMM regulation and factors responsible for TMM phenotype choice in different cells. Currently, many studies address the involvement of few genes in TMMs, but a consensus unified picture of the full process is missing.

We have developed a computational biology framework combining knowledge- and data-driven approaches to aid in understanding of TMMs. It is based on a greedy algorithm with three core modules: (1) knowledge-based construction/modification of molecular pathways for telomerase-dependent and alternative TMMs, (2) coupled with gene expression data-based validation with an in-house pathway signal flow (PSF) algorithm, and (3) iteration of these two coupled steps until converging at pathway topologies that best reflect state of the art knowledge and are in maximum accordance with the data. We have used gene expression data derived from cell lines and tumor tissues and have performed extensive literature search and multiple cycles of greedy iterations until reaching TMM assessment accuracy of 100% and 77%, respectively.

Availability of TMM pathways that best reflect recent knowledge and data will facilitate better understanding of TMM processes. As novel experimental findings in TMM biology emerge, and new datasets are generated, our approach may be used to further expand/improve the pathways, possibly allowing for making distinctions not only between telomerase-dependent and ALT TMMs, but also among their different subtypes. Moreover, this method may be used for assessment of TMM phenotypes from gene expression data, which is crucial for studies where experimental detection of TMM states is missing. Furthermore, it can also be used to assess TMM activities in proliferating healthy cells.

## Introduction

Telomeres are the ends of eukaryotic chromosomes that perform protective functions by limiting access of DNA damage response proteins to the chromosome ends, which would otherwise lead to end-to-end fusions and chromosomal instability. Telomeres get shorter after each round of DNA replication, therefore cell divisions may ultimately result in loss of proper telomeric ends [1–3]. Highly proliferative cells, such as cancer cells and stem cells have mechanisms for preserving the telomere ends after many rounds of divisions [4]. These mechanisms are collectively known as telomere maintenance mechanisms, or TMMs [5].

Currently, it is thought that there are two main TMMs: one is realized through catalytic activity of the telomerase ribonucleoprotein complex [5–7], the other occurs via homologous recombination events between telomeric sequences, and is called alternative lengthening of telomeres, or ALT [8–11]. The first mechanism is mainly employed by stem cells and many cancer cells. ALT has been identified in a fewer number of cancers. The type of TMM activated in a tumor largely determines its pathomechanistic characteristics and aggressiveness. A lot is known about the TMM mechanisms, however there is no unified picture describing the overall processes, and little is known about what leads to activation of either of the two TMM types in different cancers. Accordingly, there are no molecular pathways referring to the two processes, and in this study we’ve constructed an algorithm for reconstruction of TMM pathways according to current knowledge. Below, we have reviewed the literature to describe the known players in the two TMMs.

### Telomerase dependent telomere maintenance mechanism

Telomerase is a ribonucleoprotein complex that is able to catalytically elongate the ends of chromosomes. It is composed of the core catalytic subunit hTERT (encoded by *TERT*), the dyskerin protein (encoded by *DKC1*), and the RNA-component hTR (encoded by *TERC*), which contains telomeric template sequence [12]. While *TERC* and *DKC1* genes are ubiquitously expressed in somatic cells, *TERT* is often silenced [13]. Therefore, mere expression of *TERT* is often (but not always) enough to induce telomerase complex formation and, thus hTERT is considered as the main limiting component for telomerase assembly [13]. However, hTERT also possesses a number of extra-telomeric functions, such as regulation of gene expression, apoptosis, proliferation, cellular signaling, DNA damage response, etc., and its expression is not always correlated with telomerase activity and telomere length maintenance [14]. In this sense, study of the whole pathway leading to assembly of the telomerase complex, and its catalytic activation is of considerable value.

The *TERC* RNA component of telomerase, in humans known as hTR, is of 451 nucleotides in length and contains a telomeric template domain, and domains for interaction with hTERT [15]. Expression of *TERC* is activated through downregulation of p53, or up-regulation of c-Myc and through the action of sex and growth hormones [13]. A large part of *TERC* transcripts undergo degradation by nuclear and cytoplasmic exosomes, and a small part is processed to maturity and carried to the locations of telomerase complex assembly [16].

Dyskerin is a telomerase subunit that binds to hTR and stabilizes the telomerase complex. Mutations in the gene *DKC1* lead to a premature aging syndrome called dyskeratosis congenital. Aside from its role in telomerase complex assembly, dyskerin possesses a number of other roles, such as rRNA processing and ribosome production [17].

It’s important to note that while the presence of these three core components is obligatory for telomerase complex assembly, there may be many forms of telomerase, as other accessory proteins, such as WD40 Repeat-Containing Protein (*WRAP53/TCAB1*), pontin (*RUVBL1*) and reptin (*RUVBL2*), also participate in its formation and are sometimes considered to be parts of the complex [18, 19].

#### Expression of hTERT and its nuclear import

Expression of hTERT is regulated by multiple factors, including transcriptional activators and repressors, promoter mutations, epigenetic modifications [20] and interactions with telomeric regions of chromosome 5 [21]. Among the known transcriptional activators of hTERT are c-Myc, NF-κB, STAT and Pax proteins, and the estrogen receptor [20]. There are also transcriptional repressors, such as CTCF, E2F1, and hormone nuclear receptors [20]. Hypo- or hyper-acetylation of H3 and H4 histone marks regulate silencing and activation of *TERT* transcription, respectively, and DNA methylation at the *TERT* promoter is known to play a complex regulatory role [20]. Differentiation of pluripotent stem cells is usually accompanied with epigenetic modifications at the *TERT* promoter to induce its silencing, while the opposite process may lead to *TERT* overexpression in cancers [20]. Overall, the regulation of transcriptional activity of *TERT* is quite complex, with multiple players acting at different regulatory levels.

While transcriptional activity of *TERT* is of crucial importance, many events should follow it to ensure proper assembly and activity of the telomerase complex. For example, nuclear import of hTERT, it’s recruitment to the place of complex assembly, proper post-translational modifications and formation of a correct conformational state, as well as availability of the rest of the subunits are necessary for the complex to form, while certain post-translational modifications ensure enzymatic activity of hTERT in the already assembled holoenzyme [19].

At the first place, hTERT should be recruited to the nucleus after translation. hTERT is a large protein (^~^124 kDa), and thus it gets into the nucleus via active transport, through its bipartite nuclear localization signal (NLS) located at residues 220-240. The NLS sequence is recognized by importin-α (e.g. *KPNA1*), which regulate the process of import [22]. Akt-mediated phosphorylation of S227 of hTERT leads to efficient binding of importin-α 5 to the NLS, and promotes its nuclear import [22]. Well known upstream activators of Akt are phosphoinositide-dependent kinase-1 (PDK-1) [23], phosphoinositide 3-kinase (PI3K), as well as heat shock protein 90 (Hsp90), a well-known repressor is protein phosphatase 2A (PP2A) [24]. Of the nuclear pore complexes, Nup358 (*RANBP2*) is known to play a major role in hTERT import [25]. It’s also thought that importin 7 (*IPO7*) may mediate hTERT nuclear localization, possibly by an alternative nuclear import pathway [25]. Molecular chaperons Hsp90 and p23 (*PTGES3*) bind to hTERT and maintain its proper conformation for the nuclear import to take place [26]. In contrast to this, binding of CHIP ubiquitin ligase (*STUB1*) and Hsp70 (*HSPA1A*) leads to cytoplasmic accumulation and degradation of hTERT [22]. Another kinase that activates telomerase via hTERT phosphorylation is protein kinase C (PKC). The study by Chang *et al* [27] suggests that PKC-mediated phosphorylation of hTERT promotes its binding to Hsp90, therefore leading to proper telomerase complex assembly.

Certain proteins may also perform posttranslational modifications of hTERT that can either inhibit its nuclear import, or complex assembly, or enzymatic activity. An example is c-Abl protein tyrosine kinase, which is known to inhibit the activity of telomerase by phosphorylation of hTERT. However, the exact role of this modification is not yet understood [28].

Aside from the NLS signal, hTERT also possesses a nuclear export signal (NES). Therefore, hTERT can shuttle in and out of the nucleus. In *yeast*, hTERT is exported from the nucleus to the cytoplasm, where the assembly of the telomerase complex takes place. In humans, it is not yet known whether the assembly happens in the nucleus or in the cytoplasm. Studies have shown that mutations in the NLS signal decrease telomerase activity, but the effect of NES mutations still has to be investigated [19].

#### *TERC* transcription, processing and degradation

In contrast to *TERT*, *TERC* is not considered as a rate-limiting factor for telomerase activity, as it is relatively constantly expressed. However, when *TERT* becomes overexpressed, hTR levels may limit the amount of assembled complexes [29]. This has been demonstrated in a series of experiments in mice, where it was shown that *TERT* overexpression in the absence of *TERC* does not lead to telomerase activity, and even inhibits tumorigenesis [30].

The rate of *TERC* transcription *per se* is not enough to ensure required amounts of hTR available for telomerase assembly: one should take into account that the amount of mature hTRs depends on the competition between the processes of maturation and degradation of newly synthesized *TERC* transcripts. These processes are controlled by nuclear and cytoplasmic exosomes that degrade those nascent hTR transcripts that were marked by certain modifiers immediately after transcription; or by other modifiers that remove degradation marks and promote the export of hTRs to Cajal bodies, where they become mature [16]. Among hTR processing complexes are the CBCA complex (*NCBP1, SRRT, NCBP2*) which leads to 5’ capping, and the NEXT complex (*RBM7, ZCCHC8, MR4*) that adds oligo-A tails to hTRs [16]. CBCA and NEXT associate with hTR co-transcriptionally and form the CBCN complex, which recruits the nucleolar exosome [16]. Long hTR transcripts get degraded by the exosome (largely by its EXOSC10 component), while the short ones are transported to the nucleolus and partly become oligo-adenylated by the TRAMP complex (*MTR4, ZCCHC7 and PAPD5*) [16]. Oligo-adenylation also marks these transcripts for degradation by the exosome. A part of these transcripts escape the exosomal degradation due to the action of PARN, which removes oligoA tails from the hTRs [31]. Polymerases may add polyA tails to the hTRs, to which PABPN1 will bind to promote PARN-directed hTR maturation [16]. The mature hTRs that are 451 nts in length are then properly folded and bound by dyskerin-NOP10-NHP2 complex and NAF1, which provides final protection against degradation [19]. This complex is recruited to Cajal bodies, where NAF1 gets replaced by GAR1 [19]. Besides nuclear exosomes, cytoplasmic exosomes DCP2 and XRN1 are also factors that may degrade hTR after its export, particularly when dyskerin binding to hTR in nucleus is compromised [31]. A recent finding also highlights the role of RNA-binding protein FXR1 in stabilization of TERC transcripts and their protection from degradation [32].

#### Telomerase assembly

The assembly of telomerase is aided by ATPases pontin and reptin (encoded by *RUVB1* and *RUVB2*) [18]. It has been shown that pontin and reptin affect hTR levels through their interaction with dyskerin, however, it has not been shown if they also interact with hTERT directly. In yeast the site of telomerase assembly is in the cytoplasm, in humans, however, the site is not yet identified [19]. It has been observed that telomerase localizes to Cajal bodies (CB) for most of the cell cycle, which is driven by interactions of WRAP53/TCAB1 with hTR, but the role of CBs in complex assembly is not yet clear [33].

#### Recruitment to telomeres and catalytic activity

The localization of the assembled nucleoprotein complex to telomeres is further guided by interactions with shelterin proteins POT1, TPP1, TRF1 and TRF2 [18, 19]. An important role in recruitment of telomerase to telomeres is played by WRAP53/TCAB1, however it is not yet established whether WRAP53 mediates interaction of telomerase with telomeres, or it induces posttranslational modifications that promote telomerase-telomere interactions [19].

The catalytic activity of the holoenzyme is also regulated by shelterin proteins [34]. For example, TRF1 inhibits telomerase activity, and the longer the telomeres, the more of TRF1 is recruited, and thus the stronger will be the inhibition. Another shelterin protein, POT1, competes with telomerase by binding to the single stranded 3’ telomere overhang. Similarly, other shelterin proteins, such as TIN2, TPP1 and TNF2 also act as negative regulators. Partly due to this regulatory mechanism, and for some yet undefined factors, telomerase elongates mainly the shortest telomeres and the mean telomere length of the cells expressing telomerase usually stays at a relatively constant level [35].

Taken together, the regulation of telomerase dependent TMM occurs at multiple levels of cellular activity: starting from the expression of the enzyme components, their proper processing and maturation, and ending with proper complex assembly, recruitment to telomeres and catalytic activity. While this process is well characterized in *yeast* and other model organisms, its many aspects are still unknown in humans.

### Alternative lengthening of telomeres

A part of cancer cells keep their proliferative potential via a telomerase-independent mechanism, which depends on homologous recombination events, and is called alternative lengthening of telomeres (ALT). ALT is observed in many common tumors, but is mainly inherent to tumors of mesenchymal origin [36]. Activation of ALT is an indication of aggressiveness of tumors and poor prognosis [37]. Additionally, tumor cells may switch from telomerase positive to ALT phenotypes during development [38], and these two TMMs may also coexist in the same cell [39]. As mentioned above, there are attempts to target cancers with telomerase inhibiting therapies. In these cases, tumor cells often switch from telomerase positive to ALT phenotype [40]. The mechanisms leading to ALT activation are therefore of considerable interest, however, in spite of the significance of the topic, the existing knowledge is yet scarce. Below we describe the state of the art on known ALT mechanisms and the factors leading to their activation.

#### Descriptors of ALT phenotype

ALT positive cells are described by a number of phenotypic markers. They contain extrachromosomal telomeric repeat containing DNA, such as T-circles that are generated via resolution of telomeric T-loops by recombination enzymes, and C- and G-circles, which are largely single stranded circular sequences of C-terminal or G-terminal telomeric repeats, with C-circles being more abundant. ALT cells usually have longer mean telomere lengths: about 20 kb on average, as opposed to telomerase positive cells that usually have less than 10 kb of mean telomere length [41]. The telomere lengths at individual chromosomes of ALT cells are highly heterogeneous with simultaneous presence of extremely short and extremely long telomeres, and the length of individual telomeres rapidly changes during proliferation. In ALT cells, there is a greatly elevated level of recombination events at telomeres, and, finally, a large number of PML (promyelocytic leukaemia) bodies containing telomeric chromatin, also called ALT associated PML bodies (APBs). The role of PML bodies is not well understood, but it is known that these are involved in senescence and DNA damage response (DDR). APBs are highly characteristic to ALT cells, and can be observed both attached to the chromosome ends, and to extrachromosomal telomeric sequences. APBs contain recombination proteins and it is assumed that ALT associated recombination events happen at APB sites [42]. Therefore, presence of APBs, as well as telomeric C-circles are used as markers for identification of the ALT phenotype [43].

Additionally, it is known that the majority of ALT cells have mutations in *ATRX* and/or *DAXX* genes; however, their role in ALT initiation is not yet clearly stated [44]. Finally, telomerase is usually under-expressed in ALT cells, however the functional consequences of its down-regulation are also not known.

#### (Possible) mechanisms of homologous recombination in ALT

There is a growing experimental knowledge highlighting the sequence of events happening in ALT and the main players involved in each event. A comprehensive review by Cesare *et al* [45] has summarized the existing knowledge on ALT regulators, and the paper by Pickett *et al* [46] has discussed involvement of these factors in particular stages of homologous recombination (HR) happening in ALT.

Two possible mechanisms of telomere elongation in ALT cells are currently considered: unequal telomeric sister chromatid exchange (T-SCE) and HR dependent DNA replication. T-SCE events are generally observed with increased frequency in ALT cells and the hypothesis assumes that due to unequal length distribution of telomeres on sister chromatids, one of the daughters receives chromosomes with long telomeres and the other one with short telomeres, and thus, the first cell will gain proliferative capacity as a result of uneven segregation of the longer telomeres. The existence of a mechanism leading to such non-random segregation of long and short telomeres into separate daughter cells is still hypothetical [45]. The second mechanism referred to as HR dependent DNA replication is thought to be employed through break induced replication (BIR). In this case, the 3’ G-strand hybridizes with another telomeric C-strand of a template sequence. The latter may be telomeric end of another chromosome, a sister chromatid, an extra-chromosomal telomeric sequence (C-circles), as well as a T-loop. The G strand is elongated up to the end of the template C strand, and afterwards the strands are separated [45]. Of these two possible mechanisms, we will further refer to the ALT pathway in reference to the HR-dependent mechanism, since it’s been studied most extensively.

The BIR mechanism can occur intra-chromosomally, inter-chromosomally, or extra-chromosomally, depending on the source of the telomeric template [47]. Strand invasion leads to formation of a Holiday Junction (HJ), and replication mediated elongation of the invaded strand. The HJ is then dissolved, and the C strand of the short telomeres is elongated using the newly copied G-strand [46]. It is also thought that extra-chromosomal telomeric repeats, such as C-loops and T-loops may serve as simple templates for strand elongation [48]. Moreover, the T-loop of the same chromosome may be used as a template in a rolling-based replication [48]. Finally, the other possibility for intra-chromosomal copying may occur through looping of the G-strand and its simple replication based on the C-strand [49]. It is subject for further investigation to reveal which of these mechanisms takes place in ALT and to which extent. Here, all further discussions will be based on consideration of the classical inter-chromosomal BIR mechanism. It is worth noting that the inter-chromosomal BIR that may occur through telomere copying from a sister chromatid assumes that sister chromatids may have varying telomere lengths. There are some studies that discuss this possibility as well [49].

HR is a complex multistep process that involves interactions between various proteins participating in strand invasion, template directed synthesis and resolution of recombination intermediates.

The first step in the overall process is strand invasion, which lead to formation of an HR specific structure known as Holiday Junction. The long-range movement of the telomeric G-strand to a homologous or non-homologous chromosome required for inter-chromosomal copying may be conditioned by Hop2-Mdn1 heterodimer interaction with RAD51 [50]. The ATR and Chk1 lead to recruitment of Hop2 to the telomeres [51]. Noteworthy, RAD51 independent mechanisms have also been observed in yeast, and such a possibility is discussed in humans as well [51]. The telomeric DNA is protected from such invasions via shelterin proteins, particularly POT1, which binds to ssDNA at telomeres. RAD51 is a known promoter of DNA invasion in HR, and it is thought that POT1 should be replaced by RAD51, and that an intermediate step in this process is the loading with replication protein A (RPA) complex. The heterogeneous nuclear ribonucleoprotein A1 (hnRNPA1) has been shown to inhibit the replacement of POT1 with RPA. RAD52 is a positive regulator of RAD51 loading [46].

The second step in HR events is the strand directed synthesis. A recent paper by Dilley *et al* [51] reports that RFC1-mediated PCNA loading at the telomeric breaks recruits the polymerase 5 through its POLD3 subunit, and RFC1-PCNA is thought to be the initial sensor of telomere damage. They have shown that POLD3 is critical for break induced telomere synthesis in the majority of ALT cells. Worth to mention, the study has also shown that the break induced telomere synthesis is not dependent on the ATR induced damage signaling and RAD51. ATR-Chk1 signaling, thus, regulates telomere integrity and ALT cell survival, while RFC1-PCNA-POLD3 independently participate in telomere synthesis [51].

After the synthesis phase the Holiday Junctions and HR intermediates are dissolved by the BTR complex composed of TOP3A, BLM, RMI1and RMI2 [46]. Several proteins, such as SLX4/1 and MUS81-EME1 complexes and GEN1, may inhibit this process via resolution of HR intermediates. It seems that dissolution happens mostly after telomere synthesis, while resolution of HR intermediates interferes with telomere synthesis thereby inhibiting ALT mediated telomere lengthening [46].

#### ALT-specific heterochromatic states

Regulation of the ALT phenotype strictly dependents on chromatin structure and the compact state of telomeres. Lack of shelterin proteins and telomeric histone modifiers leads to decompaction of telomeres and formation of an ‘open’ chromatin state. It is assumed that this open state recruits recombination proteins to the accessible telomeric DNA, which is the main potent trigger of ALT phenotype establishment [52, 53]. Particularly, all the experiments with impaired telomeric chromatin, have observed large numbers of APB bodies [53–57].

According to this notion, the proteins found to be regulating (or participating in) ALT are functionally related to one or many of the following processes: loosening of heterochromatin (histone (de)acetylases/(de)methylases, NuRD-ZNF827 complex) [58], T-loop breakage through replacement of shelterin proteins with ALT initiators (POT1, RPA, RAD51) [46], homologous-recombination (SLX4-SLX1, MUS81-EME1, BTR complex) [46], and APB formation (MRN, SMC5/SMC6 complexes, BRCA1, BLM) [59, 60].

It has been suggested that nuclear receptors NR2C2 (TR4) and NR2F2 (COUP-TF2) bind to telomeric C-type variants and recruit a recently discovered protein ZNF827 [58]. ZNF827 in turn recruits the NuRD complex to telomeres to promote ALT via increased T-SCEs, APBs and C-circles. A number of proposed mechanisms may act to support the ALT-promoting activity of the NuRD-ZNF827 complex. First, it’s thought that NuRD is able to simultaneously interact with several ZNF827 proteins. If these proteins are located at telomeres of different chromosomes, this will lead to so called “telomeric bridge” formation, which in turn may promote telomeric strand migration as an initiator of HR at telomeres. Second, the NuRD-ZNF827 complex may recruit HR proteins, such as BRIT1 and BRCA1 to telomeres [58]. Finally, the hystone deacetylases HDAC1 and HDAC2 that are part of the NuRD complex may contribute to deacetylation and reduced chromatin compaction at telomeres [58].

The NuRD complex consists of six functional subunits, and performs a number of roles in regulation of replication, transcription and genomic stability. Depending on the protein composition of the subunits, NuRD may perform specific functions, which may have opposing effects, such as tumor suppressive or promoting [61]. It is not established whether the composition of NuRD subunits affects recruitment by the ZNF827 complex. Of note, the expression of some NuRD components, as reported in [58] is not significantly different in ALT positive and negative phenotypes, leading to the assumption that the recruitment of NuRD to telomeric sites affects ALT more (if not at all), than the general abundance of NuRD proteins.

It has also been demonstrated that depletion of chromatin modifiers SUV39H1/H2, and SUV420H1/H2 which deposit the repressive H3K9me3 and H4K20me3 marks respectively, leads to chromatin decompaction, and formation of an ALT permissive environment [57, 62].

#### APB bodies

Promyeolocytic leukemia bodies (PML) are generally present in all somatic cells, and they grow in size and number as the cell undergoes senescence [63]. The PML bodies that are associated with telomeric sequences are frequently found in ALT cells (thus, the name: ALT associate PML bodies (APB)), and are considered to be the main site where HR events occur [42]. Owing to the increased number of APB bodies in ALT cells, those have been used as markers of ALT activity [43]. APB bodies are found near telomeres at chromosome ends, as well as extra-chromosomal telomeric repeats. In addition to the regular components of PML bodies, such as the PML, Sp100 and shelterin proteins, APBs also contain additional components, such as RAD1, RAD9, RAD51, RAD52, RPA, RAD51D, BLM, WRN, RAP, BRCA1, MRE11, RAD50, and NBS1, *etc* [59]. While some of the APB components participate in HR events, the role of others in ALT is not known. These are the SMC5/6 complex (NSMCE2 (MMS21), SMC5, SMC6) and the MRN complex (NBN, RAD50 and MRE11A). The MRN complex appears in the early stages of dsDNA repair, and functions as a regulator of cell cycle checkpoints. In ALT cells, the MRN complex colocalizes with HR proteins, and depletion of this complex leads to reduced telomere length [60]. Overexpression of SP100 sequesters the MRN complex away from APBs and inhibits ALT [59]. The SMC5/6 complex proteins sumoylate telomere binding proteins such as TRF1 and TRF2, leading to increased APB formation at telomeres. Inhibition of SMC5/6 complex suppresses HR at telomeres [64]. The RecQ helicase WRN is found at APBs, however, its role in ALT is not established, since it’s shown to be required in some, but not all of the ALT cells [65].

#### The role of ATRX and DAXX

The majority of ALT cells have loss-of-function mutations in either component of the ATRX/DAXX complex. There are several hypothesis of how ATRX and DAXX affect ALT, however their exact role has not been established yet [66]. The ATRX/DAXX complex is shown to deposit the histone H3.3 variant at telomeres, which may lead to repressed telomeric transcription and reduction of TERRA transcripts [67]. The role of H3.3 variant at telomeres is not clearly established, however association of ATRX/DAXX with H3.3 at telomeres has been shown to have stabilizing effect on telomeric heterochromatin, since the ATRX/DAXX complex possesses chromatin remodeling activities [68]. Moreover, in mice, ATRX has been shown to reduce HP1α and lead to chromatin decompaction [66]. Finally, a recent study has shown that preferential elongation of overhangs at the lagging chromatids promotes ALT, and ATRX was shown to partially suppress this process [69]. Despite the frequent mutation rate of ATRX/DAXX, its worth to mention that their mutations alone are not enough to initiate ALT, as has been shown for SV-40 hTERT immortalized cell lines [44].

#### Telomere fragility and sister chromatid loss

Flap structure specific endonuclease 1 (FEN1) is a canonical Okazaki fragment processing protein. Recently it has been shown that it plays a major role in telomere stabilization during replication, both at the leading, as well as the lagging strands [70]. At the lagging strand, FEN1 participates in replication fork progression, reinitiation and telomere stability [70]. At the leading strand, FEN1 cleaves the RNA:DNA hybrids that are generated during TERRA transcription. Removal of RNA fragments from the DNA as the replication progresses is important to limit telomere fragility and promote DNA repair at the leading strand [70]. Owing to its ability to stabilize telomeres, FEN1 may be important for telomere lengthening in ALT cells. However, there is a controversy, since FEN1 inhibits TERRA RNA:DNA hybrid formation, and these hybrids are known to support recombination events in ALT [70]. Studies have revealed that depletion of FEN1 leads to sister chromatid loss in ALT telomeres. It has been shown that FEN1 stabilizes telomeres in ALT positive cells. Its depletion from ALT positive cells leads to generation of telomere dysfunction induced loci (TIF) and end-to-end chromosome fusions. In telomerase positive cells, loss of telomeres at sister chromatids caused by FEN1 depletion was compensated by the action of telomerase [71].

Taken together, the mechanism through which ALT takes plays may vary, depending on the source of the template telomeric sequence and the mode of replication. Numerous studies have tried to reveal the molecular factors involved in ALT events, and these pieces of puzzle rapidly come together. It is possible that many pathways exist that lead to ALT induction and realization, however which of them is targeted by this or that study is largely not known. Therefore, currently accumulated knowledge on ALT may reveal a generalized picture of all the possible mechanisms that lead to recombination driven telomere elongation.

#### Problem statement

The studies addressing TMMs are performed at a low-throughput level: each study attempts to investigate the role of one or two molecular factors in these processes. A couple of studies have attempted to use standard data mining and functional annotation pipelines for investigation of TMMs. The main shortcoming of these studies is that there are currently no properly annotated functional categories and no computational model or framework that could validate computational means of TMM studies. Partly, this is explained by knowledge gaps in understanding of the factors that lead to activation of the TMMs, as well as the conflicting results obtained by independent studies on TM mechanisms. Additionally, the studies targeted at understanding the role of various factors in TMMs are paying attention to individual genes/proteins, and do not summarize the overall process. Moreover, data driven approaches that attempt to analyze high-throughput data to obtain lists or networks of genes associated with TMMs, fail to provide biologically meaningful consistent results.

In this paper we have implemented a methodology for construction and modeling of signaling pathways that reflect the recently accumulated knowledge on TMM related events, and that are in maximum accordance with gene expression data.

## Results

### The framework for TMM pathway reconstruction and testing

We have developed a computational biology framework combining knowledge- and data-driven approaches to aid in our understanding of the processes underlying telomerase dependent and alternative TMMs. Our framework is based on a greedy algorithm with three core modules: (1) knowledge-based reconstruction/modification of molecular pathways for telomerase-dependent and alternative TMMs, (2) coupled with gene expression data-based validation, and (3) iteration of these two coupled steps until converging at pathway topologies that best reflect state of the art knowledge and are able to make the best distinction between ALT positive and Telomerase positive phenotypic states based on available gene expression data.

1. Telomerase- and ALT TMM pathways represent graphs, where nodes are genes/proteins/RNAs and edges are physical interactions and functional effects of type activation/inhibition. The involvement of each node and interaction is derived from literature.
2. At each step of our greedy algorithm, a novel node or an interaction change is introduced in the pathway and validation is performed by computing the value of a fitness function. The fitness function is computed with an in-house pathway signal flow algorithm (PSF) that estimates the activity of telomerase- and ALT- TMM pathways based on gene expression values and signal propagation through pairwise interactions between nodes. Based on PSF activity values of the two pathways it is possible to assess which TMM (if any) is active in different samples. The fitness function is computed by comparison of the computational PSF assessments with experimental results on TMM phenotype detection and consists of two types of metrics:

a. Classification accuracy based on 2-class SVM-based separation of ALT positive and ALT negative, as well as Telomerase positive and Telomerase negative samples;
b. Median difference of PSF values for ALT and Telomerase pathways between different phenotypic groups.
3. Decision making: if the modification increases the fitness function value, it is kept in the pathways, and is discarded otherwise. The steps 1-3 are repeated until all the literature-derived gene-players are tested and until relative convergence of the fitness function values.

### Reconstruction of the TMM pathways

Using the above mentioned algorithm and around hundred publications on the mechanisms of activation and realization of Telomerase and ALT TMMs, we have constructed several versions of ALT and Telomerase pathways. We have started with an initial pathway with well documented members and interactions, and iteratively added nodes and edited their positions in the pathways to arrive at the best classification power. Figure 1 shows the changes in prediction accuracy metrics with each pathway extension step for cell lines and for liposarcoma tumors. The classification accuracy increases for both datasets during iterations, and achieves the highest points at the last iterations. The rest of the fitness function metrics – PSF median differences between different phenotypic groups for ALT and Telomerase pathways – are varying, dynamically converging at an optimal level.

**Figure 1.**
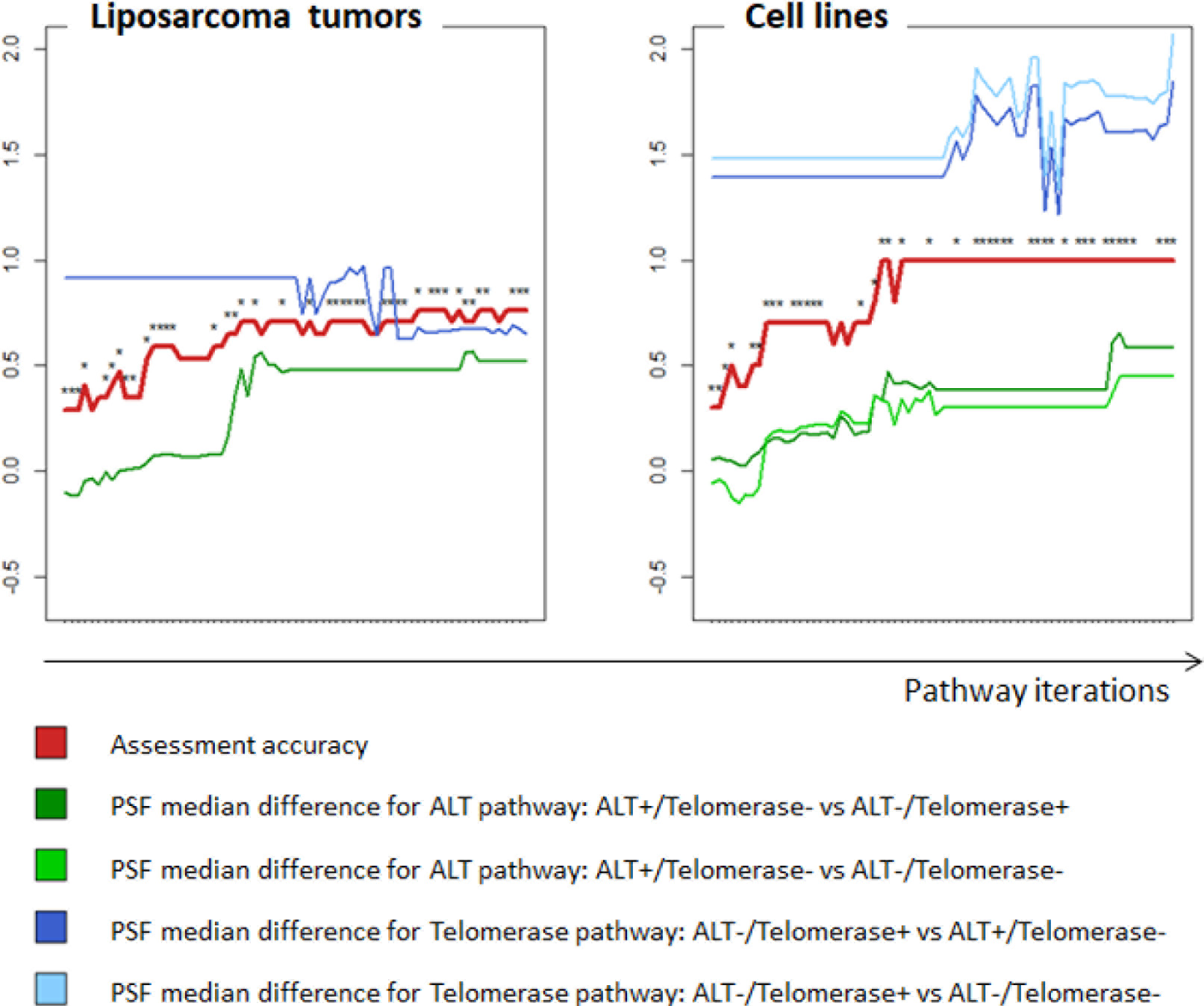
Assessment accuracy dynamics of TMM pathways during pathway extensions. The two plots represent TMM pathways’ fitness function dynamics during pathway modifications for liposarcoma tumors (**left**) and cell lines (**right**) respectively. The x-axis shows pathway iterations: either the Telomerase or ALT pathway is modified at each iteration step. Performance scores are represented on the y axis. The red lines correspond to the assessment accuracy changes, as predicted by SVMs (see methods, Pathway performance assessment). The dark green and light green lines represent the average difference of the ALT pathway activity between ALT^+^/Telomerase^−^ and Telomerase^+^/ALT^−^ samples (dark green); and between ALT^+^/Telomerase^−^ and ALT^−^/telomerase^−^ samples (light green). The dark blue and light blue lines show the average difference of the Telomerase pathway activity between Telomerase^+^/ALT^−^ and ALT^+^/Telomerase^−^ (dark blue); and between Telomerase^+^/ALT^−^ and ALT^−^/telomerase^−^ samples (see methods, Pathway performance assessment).

#### The telomerase pathway

The Telomerase pathway at the final iteration is depicted in Figure 2 A. The pathway consists of three main branches. The yellow branch includes the factors that lead to nuclear localization, enzymatic activation of hTERT. These are importins, nuclear pore proteins and hTERT interactors, such as p23 and heat shock protein 90. We have not included all the factors mentioned in the introduction, since not all of them were increasing the accuracy of pathway based TMM assessment algorithm. Since there are three isoforms for HSP90, we have chosen the one with maximum abundance, as depicted in the node functions.

The violet branch involves the genes and complexes involved in hTR transcription and maturation, while the green one includes the factors leading to degradation of hTR transcripts. The amount of mature hTR transcripts depends on the interplay between these two processes. It’s worth noting that only PAPD5 was included in the TRAMP complex, and only EXOSCIO was included from the exosomes – the rest were reducing the fitness function metrics.

The gray branch gathers factors leading to proper assembly of the telomerase complex, such as Ponting (*RUVB1*) and Reptin (*RUVB2*). Finally, the blue nodes highlight the core subunits of the telomerase complex and the sink node of the pathway (“Telomerase“).

**Figure 2.**
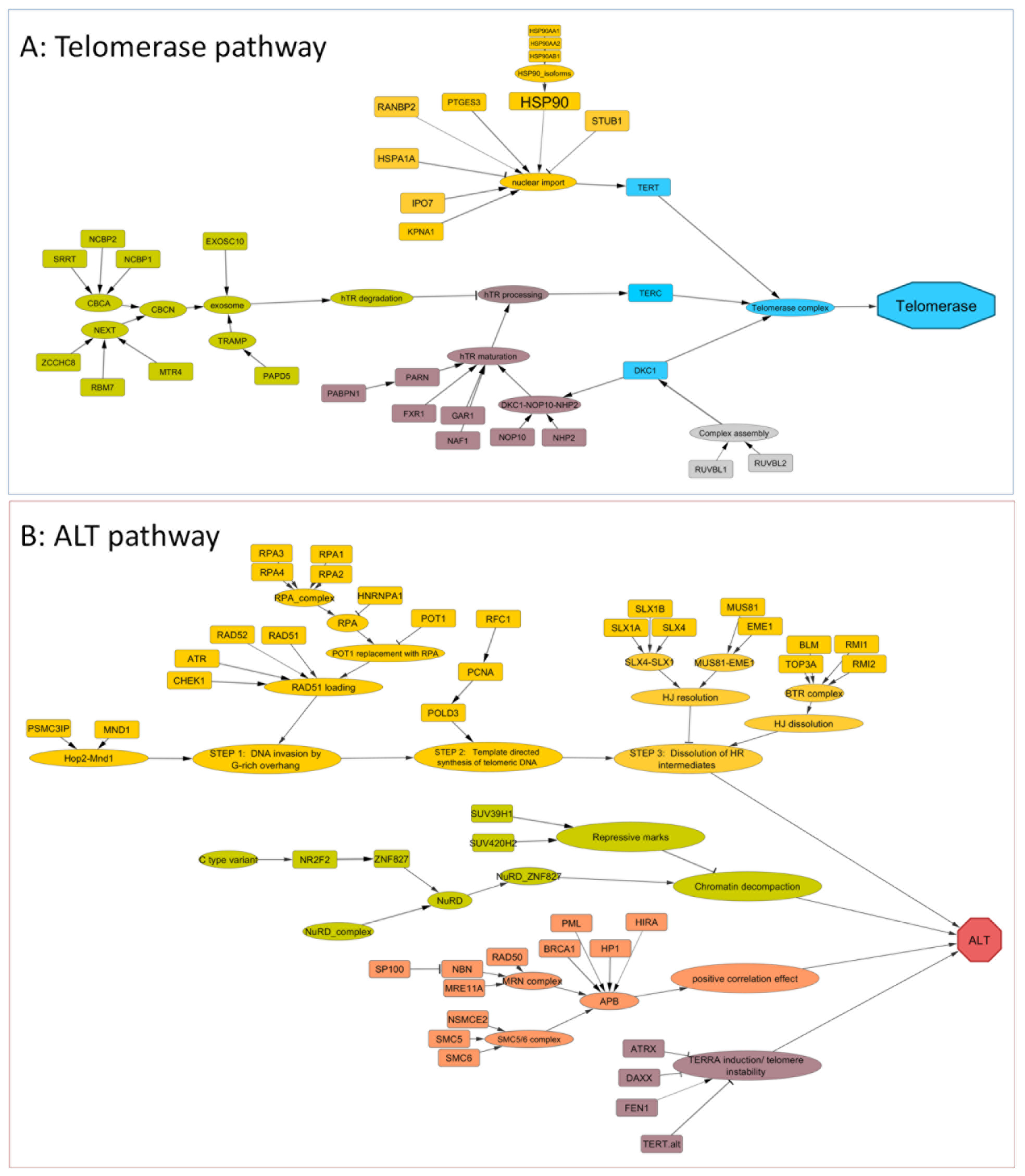
The Telomerase (A) and ALT (B) pathways. The “Telomerase” and “ALT” nodes are the targets of each pathway. In **(A)**, the core components of the telomerase complex are colored blue. The rest of the colors in both plots are for enhancement of readability. Rectangular nodes represent genes, while oval-shaped nodes represent processes or complexes. The edges with delta and T shaped targets indicate activation and inhibition interactions, respectively.

#### The ALT pathway

The ALT pathway is depicted in Figure 2 B. As is described in the Introduction, multiple mechanisms exist that allow for template directed synthesis of telomeres in ALT: the template may be from sister chromatids (intra-chromosomal), from distantly located chromosomes (inter-chromosomal) and from extra-chromosomal telomeric sequences. In construction of the ALT pathway, we have accounted for the factors involved in inter-chromosomal break-induced repair and extra-chromosomal template-driven mechanisms. It is important to note, however, that our pathway mostly represents a generalized picture, where the mechanism that is realized by involvement of this or that pathway entity is not clearly indicated.

In the ALT pathway, the yellow branch involves the main processes leading to HR in ALT. These processes are divided into three consecutive steps. The first step presented with the node “Step 1: DNA invasion by G-rich overhang” involves the factors leading to ssDNA loading with RAD51, which leads to invasion of the G-rich strand onto a telomeric template. It is worth noting that the RAD51-dependent pathway is observed in the majority, but not all of the ALT cells. The Hop2-Mnd1 heterodimer is also not an exclusive player in ALT. It plays an essential role in some ALT cells by promoting strand invasion onto distantly located chromosome ends. The second step represents the telomere strand synthesis after strand invasion, and is presented by the node “Step 2: template directed synthesis of telomeric DNA”. It’s been shown that this step is performed by DNA polymerase 5. It is reported that its POLD3 subunit, but not POLD4, is essential for this step. Indeed, inclusion of the POLD4 node did not improve the pathway performance and thus was removed. The final step is the dissolution of the Holiday Junctions after the telomere synthesis is finished (node “Step 3: Dissolution of HR intermediates“). An opposite process to this is the resolution of HR intermediates that occurs in the absence of telomere synthesis.

The rest of the branches involve factors that promote ALT by creation of an ALT permissive environment or those that are positively/negatively correlated with ALT by yet unknown mechanisms. The green branch represents the factors leading to decompaction of chromatin and establishment of ALT permissive environment, such as chromatin modifiers (the SUV family), and the NURD-ZNF827 complex. Notably, this complex has additional roles in ALT, other than chromatin decompaction, as mentioned in the introduction. The node named “C-type variants” represents telomeric TCAGGG repeats that are often found in ALT and recruit nuclear receptors. This node did not have an initial FC value in our calculations. However, if there were whole genome sequencing data along with gene expression, one could compute the abundance of C-type variants and use it in the PSF calculations. Of note is that from the two nuclear receptors mentioned in the introduction (*NR2C2*, *NR2F2*), only NR2F2 was leading to increased accuracy. Inclusion of NURD complex subunits also appeared not beneficial, as is supported by the fact the expression of the subunits *per se* is not affecting ALT activity – rather the recruitment of the subunits to APB bodies [58].

The orange branch involves proteins found in APB, and factors supporting or inhibiting APB formation.

The violet branch involves factors that are associated with heterochromatic state or stability of telemetric DNA ends. While ATRX/DAXX genes are mutated in the majority of ALT cells, the depletion of wild variants also promotes ALT activity [69]. We did not have genome variant data for the studied samples, and thus have considered higher expressions of these genes to have inhibitory effects on ALT. Availability of information on genetic variations would probably lead to more accurate results. Absence or low abundance of telomerase is also promoting ALT, since it leads to telomere shortening and instability at telomeric ends. Low *TERT* expression also usually provokes the cells to escape senescence via the alternative ALT mechanism. Finally, *FEN1* has a role in stabilization of telomeres and inhibition of sister chromatid loss. As mentioned in the Introduction, the role of FEN1 is controversial, since it also reduces the RNA:DNA hybrids that are known to promote ALT.

### Pathways’ prediction accuracy for cell lines

We have computed PSF values for ALT and Telomerase TMMs using gene expression data from four ALT^+^/Telomerase^−^ (SKLU, WI38-SV40, SUSM1, KMST6), four ALT^−^/Telomerase^+^ (5637, C33A, HT1080, A2780), two ALT^−^/Telomerase^−^ (IMR90, WI38) cell lines. The boxplots of PSF values for both TMM pathways are depicted in Figure 3. The ALT^+^/Telomerase^−^ cell lines had significantly higher PSF values compared to ALT^−^/Telomerase^−^ (median differences = 0.45) and ALT^−^ /Telomerase^+^ (median difference = 0.58) cell lines (p value = 0.002). A similar picture is observed for the Telomerase TMM, with ALT^−^/Telomerase^+^ cell lines scoring significantly higher than the ALT^−^/Telomerase^−^ (median differences = 2.07), and ALT^+^/Telomerase^−^ (median difference = 1.84) cell lines (p value = 0.002). Thus, in this case, the ALT and Telomerase^+^ TMMs alone are powerful enough to distinguish between ALT^+^ and Telomerase^+^ cell lines from other phenotypic groups (Figure 3).

The combined assessment of PSF based support vector machine classification allowed us to distinguish between the three phenotypic groups simultaneously (Figure 4).

**Figure 3.**
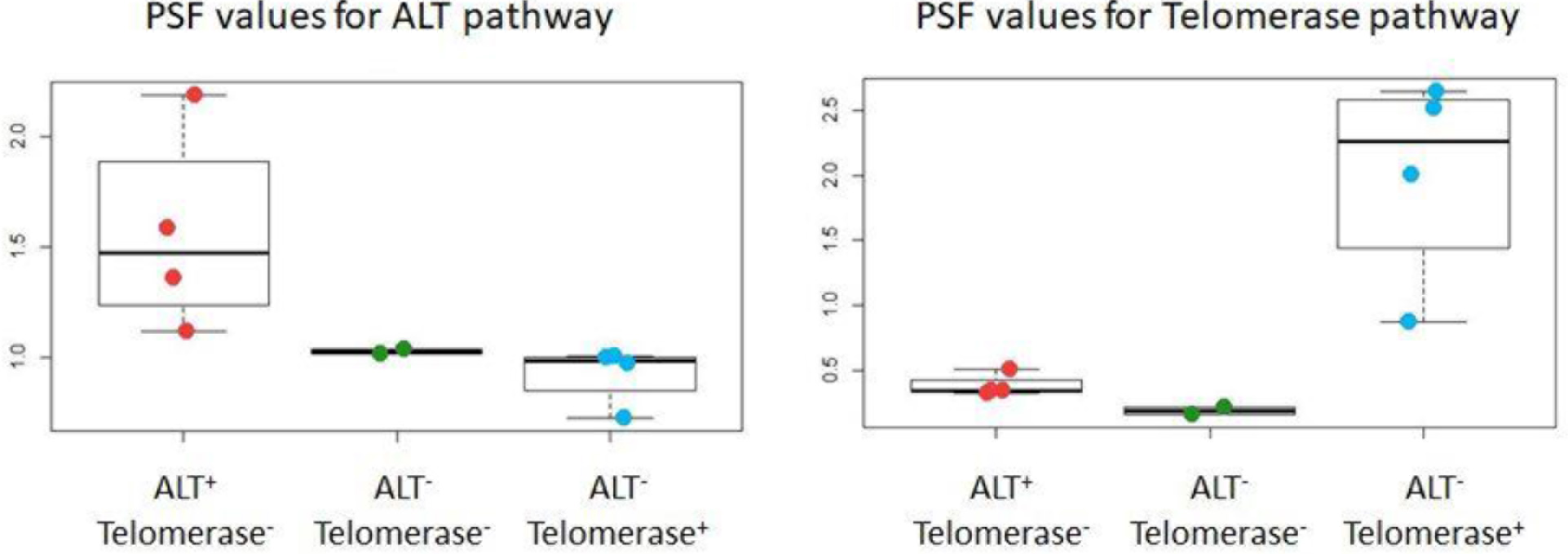
Boxplots of group-wise PSF values for each TMM for cell lines. The red, green and blue dots represent ALT^+^/Telomerase^−^, ALT^−^/Telomerase^−^ and ALT^−^/Telomerase^+^ cell lines respectively. ALT^+^/Telomerase^−^ cell lines have higher PSF values for the ALT TMM compared to the other two groups (**left**), while the Telomerase TMM shows considerably higher PSF values for ALT^−^/Telomerase^+^ cell lines (**right**). These differences are statistically significant.

**Figure 4.**
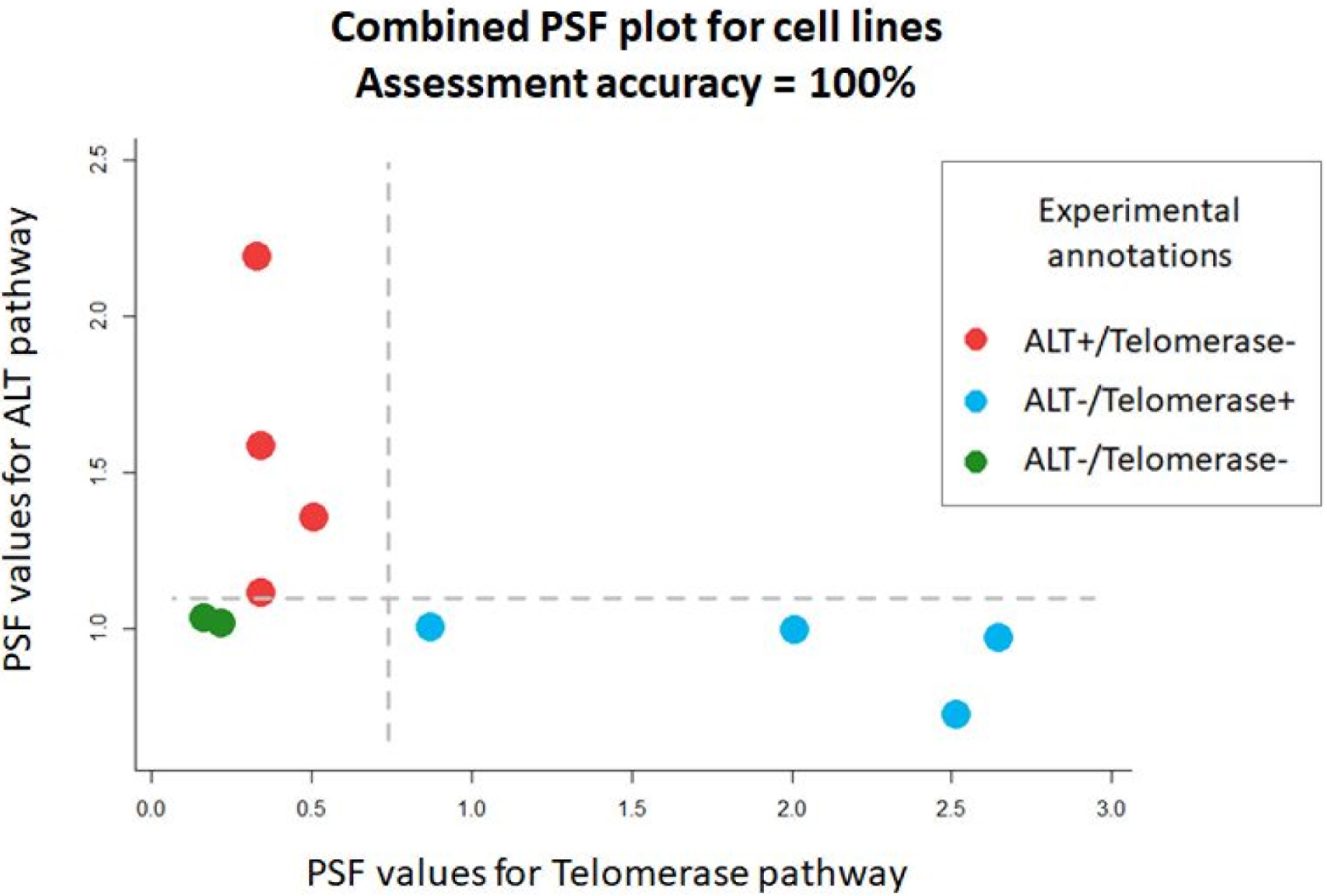
Paired PSF values for ALT and Telomerase TMMs in cell lines. The PSF values for the Telomerase TMM are on the x axis, and for the ALT TMM those are on the y axis. The red, green and blue dots correspond to ALT^+^/Telomerase^−^, ALT^−^/Telomerase^−^ and ALT^−^/Telomerase^+^ cell lines, respectively. The horizontal and vertical dotted lines separate ALT^+^ from ALT^−^, and Telomerase^+^ from telomerase^−^ phenotypes, respectively. The thresholds are determined by one-dimensional linear SVMs (see methods). The bottom-left, upper-left and bottom-right quadrants classify ALT^−^/Telomerase^−^, ALT^+^/Telomerase^−^ and ALT^−^/Telomerase^+^ phenotypes respectively. The prediction accuracy, computed by the ratio of correct versus incorrect classifications, is 1 in this case, as all the classifications correspond to the actual phenotypes.

### Pathways’ prediction accuracy for the tumors/hMSC group

The tumor group was comprised of liposarcoma samples, where nine had ALT^+^/Telomerase^−^ and eight had ALT^−^/Telomerase^+^ phenotypes. The gene expression dataset also included four hMSC samples derived from the bone marrow of healthy individuals. Since it is not clear what TMM is active (if any) in hMSCs, we have not included them during iterations. However, we have added their PSF values to the plots later, for comparison.

The PSF median differences between ALT^+^/Telomerase^−^ and ALT^−^/Telomerase^+^ tumors for ALT (median difference 0.59, p value 0.011) and Telomerase (median difference 0.67, p value 0.008) pathways were significant. The Telomerase pathway PSF scores were also significantly higher in ALT^−^/Telomerase^+^ tumors compared to hSMCs (median difference 0.95, p value 0.004). However, the ALT^+^/Telomerase^−^ tumors had only slightly higher ALT activity compared to hMSCs (median difference 0.3, p value 0.148) (Figure 5).

**Figure 5.**
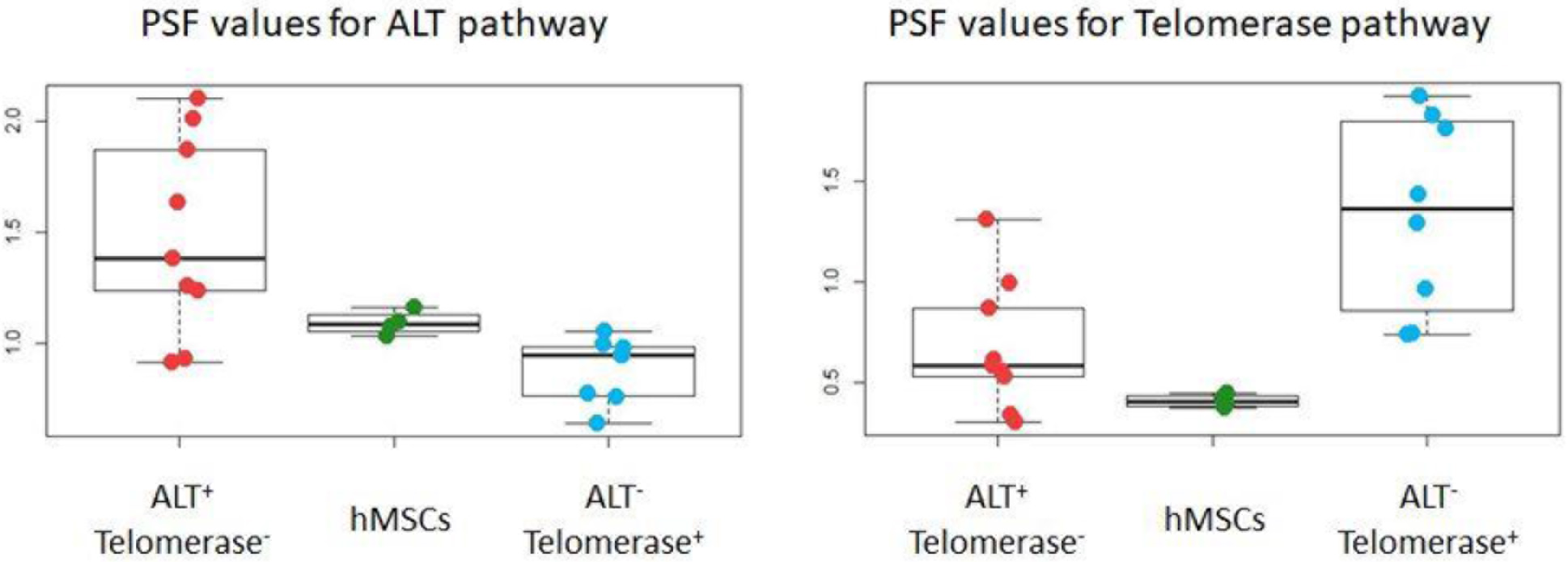
Boxplots of group-wise PSF values for each TMM pathway for liposarcoma tumors/hMSCs. The red, blue and green dots represent ALT^+^/Telomerase^−^, ALT^−^/Telomerase^+^ tumors and hMSCs, respectively. ALT^+^/Telomerase^−^ tumors have considerably higher PSF values for the ALT TMM compared to ALT^−^/Telomerase^+^ tumors (median difference 0.59, p value 0.011), and were non-significantly higher values compared to hMSCs (median difference 0.3, p value 0.148) (**left**), while the Telomerase pathway shows considerably higher PSF values for ALT^−^/Telomerase^+^ tumors compared to both ALT^+^/Telomerase^−^ tumors (median difference 0.67, p value 0.008) and hMSCs (median difference 0.95, p value 0.004) (**right**). All these difference, except for ALT^+^/Telomerase^−^ versus hMSC comparison for the ALT TMM, are statistically significant.

Combination of the ALT and Telomerase PSFs followed by SVM based classification for tumor samples only, revealed a classification accuracy of 76%, with two misclassified ALT^−^ /Telomerase^+^ samples, and two misclassified ALT^+^/Telomerase^−^ samples (Figure 6). To see where on this plot hMSCs are, we have plotted them on the right plot of Figure 6. hMSCs tend to shift more towards the ALT^−^/Telomerase^−^ quadrant, although a couple of them appear in the ALT^+^/Telomerase^−^ quadrant.

**Figure 6.**
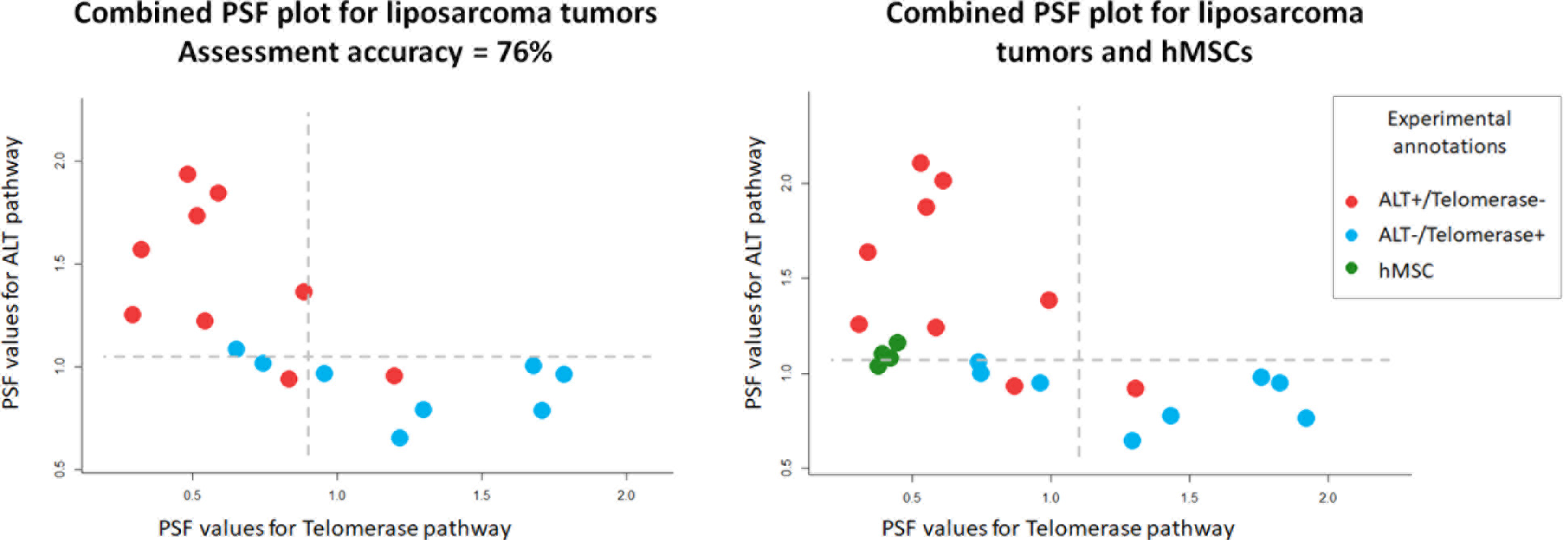
PSF values for ALT and Telomerase TMMs in liposarcomas and hMSCs on a 2D space. The PSF values for the Telomerase TMM are on the x axis, and for the ALT TMM those are on the y axis. The red, blue and green dots correspond to ALT^+^/Telomerase^−^, ALT^−^/Telomerase^+^ tumors and hMSCs, respectively. The horizontal and vertical dotted lines, determined by linear SVMs, separate ALT^+^ from ALT^−^, and Telomerase^+^ from Telomerase^−^ phenotypes, respectively. The bottom-left, upper-left and bottom-right quadrants classify ALT^−^/Telomerase^−^, ALT^+^/Telomerase^−^ and ALT^−^/Telomerase^+^ phenotypes respectively. The prediction accuracy, computed by the ratio of correct versus incorrect classifications, is 0.76 in the case of tumor samples, with four misclassified samples out of 17. hMSCs were not included in computations of assessment accuracy, since their phenotypes are not experimentally determined. They are plotted on right plot for comparison.

### Comparison to functional annotation analysis

Functional annotation using over-representation (ORA) [72] or gene set enrichment analysis (GSEA) [73] is the usual pipeline applied to understand the phenotypic differences between two groups of samples under investigation. We have performed ORA on our test datasets (cell lines and liposarcoma group) to compare the power of that methodology to our TMM based approach. For this we have conducted gene-wise Kruskal-Walllis tests to obtained genes differentially expressed in ALT^+^/Telomerase^−^ and ALT^−^/Telomerase^+^ cell lines/tumors. Multiple test correction with Benjamini-Hochberg method did not revealed any significantly deregulated gene nor in cell lines, neither in tumors. We have thus taken top 100 genes with unadjusted p values of less than 0.01, and have supplied them to functional annotation packages David (https://david.ncifcrf.gov/) [74, 75] and Webgestalt (http://webgestalt.org/) [76]. According to the results, no functional category was significantly enriched. Among the non-significantly enriched terms/categories, none was explicitly related to telomere maintenance mechanisms. Moreover, there was only one gene (*NANS*) common between the two lists of differentially expressed genes, which means that the lists are not identified by ALT^+^ versus telomerase^+^ phenotypes, but possibly by other phenotypic differences between the two sets of cells. Finally, none of the genes included in our TMM pathways was present in the lists of top 100 differentially regulated genes.

Previously, Lafferty-Whyte *et al* [77], had identified a set of 297 genes that were able to cluster ALT^+^/Telomerase^−^ and ALT^−^/Telomerase^+^ cell lines and liposarcoma samples. We have obtained the list of the genes in this signature by request from the authors, and have submitted it to overrepresentation analysis by Webgestalt. According to the results, these genes were significantly enriched in GO terms associated with Golgi vesicle transport (GO:0048193, FDR =
0. 16, 10 genes), Protein transport (GO:0015031, FDR = 0.16, 24 genes) and Establishment of protein localization (GO:0045184, FDR = 0.16, 25 genes). No category or pathway related to telomeres was enriched. Comparison of the signature set with the genes involved in our TMMs revealed three common genes: *TERT* and *PRKCA* from the Telomerase pathway, and *PRK* and *POLD3* from the ALT pathway.

### Validation of pathway signal flow algorithm

To demonstrate the added value of the pathway signal flow algorithm based classification, as opposed to gene set analysis, we have performed hierarchical clustering of the samples with the gene sets obtained from our ALT and Telomerase pathways. The results are available in supplementary material (Supplementary figures S1-4). Hierarchical clustering based on expression of the genes involved in ALT and Telomerase TMMs did not properly classify the samples, thus showing the necessity of accounting not only for the genes, but also their interactions within the pathways.

### TMM app and project website

Each iteration of the TMM pathway reconstruction algorithm consists of fold change computation from gene expression data, PSF computation on TMM pathways and statistical analysis. All these steps are performed automatically by TMM app for Cytoscape. The app is available for download from the Cytoscape App Store (http://apps.cytoscape.org/apps/tmm). We have designed a webpage for the TMM project, which contains project description, links to download the pathways, the data and the example reports, and the app user guide.

#### New pathway submissions

Researchers may modify the TMM pathways by adding/deleting genes, modifying interactions. These modifications, if grounded by better performance in terms of TMM assessment, in accordance with experimental validations, may find their place in the Network downloads section of the TMM project page. To make all the modification details available and to have a database of TMM candidate genes, we have created a table containing all the genes ever tested or those to be tested for inclusion in the TMM pathways at: http://big.sci.am/software/tmm/tmm-genes/. To submit their modifications, researchers should upload the files at the submissions section of the TMM webpage (http://big.sci.am/software/tmm/#submissions). Those will be reviewed by us to reproduce the reports and validate the changes, and will be uploaded after approval for public use.

## Discussion

Telomere maintenance and elongation mechanisms have long been investigated, and have gained particular attention in the last decade. Those are interesting to many labs focused on basic research in aging, stem cells, senescence etc., as well as to those engaged in translational research, particularly in development of TMM targeting anticancer therapies. While the TMM pathways are laborious to study experimentally, a lot of information has already been accumulated. However, the main obstacle for their efficient utilization is the absence of well-established and valid TMM models and poor utilization of the recent developments in high-throughput technologies, mainly because of the lack of respective computational methodologies.

The TMM pathways we have created here were intended to serve as a computational model to integrate the studies of telomere maintenance mechanisms into high-throughput data analysis pipelines. Gene expression-based assessment of TMM activities has allowed us to assess the TMM phenotypes of ten cell lines with 100% accuracy. The accuracy of assessing the phenotypes of liposarcoma tumors was 76%, with three misclassified ALT^+^/Telomerase^−^ tumor and one misclassified ALT^−^/Telomerase^+^ tumor. The misclassification of liposarcoma samples may be explained by several possible reasons, such as probable incompleteness of the TMM pathways, as well as the sensitivity/specificity of the experimental assays for TMM detection. In liposarcoma samples, telomerase activity has been measured by TRAP assay [78] and ALT had been detected by measuring the amount of APB bodies [78]. It is worth mentioning that, as other PML bodies, APBs may be generated because of cellular senescence, and are not necessarily indicative of ALT activity in the cell [63]. It is a question of further investigations whether the misclassified tumor samples are, in fact, misclassified by our approach, or they are misclassified by the experimental assays. On the other hand, the possibility of existence of ALT positive and telomerase positive cells within the same tumor should also not be excluded. In other words, it’s possible that RNA-sequencing be performed on ALT^+^/Telomerase^−^ cells and TMM detection – on ALT^−^/Telomerase^+^ cells derived from the same tumor, and vice versa, thus leading to the observed misclassification.

Currently, the TMM status of human mesenchymal stem cells is not established: many studies have identified very low or undetectable levels of telomerase activity [79], and some have performed ALT activity assays, but have not detected significant levels of ALT markers [80]. Therefore, we have not included those during pathway reconstruction. Later on, we have checked to which phenotype the hMSCs belong to, according to our classifier. The hMSCs tightly clustered together, and were at the edge of ALT^+^/Telomerase^−^ and ALT^−^/Telomerase^−^ quadrants (Figure 6, left plot). It is known that the majority of tumors with ALT activity originate from mesenchymal cells [36]. The fact that we have detected ALT activity in hMSCs tempts to speculate that mesenchymal cells normally have high expression levels of genes involved in the ALT pathway, which may ultimately foster these cells to more easily convert to ALT phenotype during malignant transformations.

In order to compare the power of our methodology to that of functional annotation analysis, we have used the lists of genes showing differential expression in ALT^+^ versus Telomerase^+^ samples, as well as have used the list of genes generated by the Lafferty-Whyte *et al* [77] based on regression analysis, and submitted those sets to Overrepresentation analysis (ORA). None of the lists did return any functional category associated with telomere maintenance and lengthening mechanisms. It’s important to take into consideration that larger sample sizes might produce lists more related to TMM mechanisms. However, while the ORA or GSEA approaches are valuable and easy-to-use methods for identification of functional categories able to distinguish between studied groups, they have two main disadvantages that make them invalid for use in TMM prediction or for identification of genes related to TMM. First, the lists of differentially expressed genes speak not only about the phenotype of interest, but also about other factors, such as batch effects, sampling bias, etc. Thus, functional annotation analyses on different samples with the same phenotypic difference will, in most of the cases, reveal inconsistent results. On the other hand, they strictly depend on the quality and completeness of respective functional categories. So far, there is no publicly available complete functional category/pathway that has thoroughly collected genes related to ALT or Telomerase pathways. The most relevant functional category is the “Regulation of telomere maintenance via telomere lengthening” term available in the Gene Ontology database (GO:1904356). There are 75 *Homo sapiens* gene annotations to this term. Some of those are part of our TMM pathways, however most of them are genes that regulate expression of *TERT* or other genes; some indirectly regulate telomere maintenance; and some genes encode shelterin or telomere-interacting proteins. Of important consideration is also the fact that enrichment of DEG genes in one functional category does not imply whether that category is activated or repressed in the studied samples: some of the genes in the category may be under- expressed and some may be over-expressed, which in many cases will lead to an ‘averaging out’ effect.

Overall, data driven methods, such as functional annotation clustering, are powerful tools to detect unknown differences between two groups of samples. However, functional annotation analysis is not powerful in the particular case of detecting the TMM phenotypes, because of the lack of properly curated and annotated functional categories or pathways. Another shortcoming of data driven methods is their inconsistency across studies. Mining for differentially regulated genes between ALT^+^ and Telomerase^+^ samples will reveal varying sets of genes, as was the case in [77]. Moreover, the genes associated with the TMM phenotypes revealed by data driven methods are not necessarily involved in TMM activation pathways: they may have regulatory functions, indirect effects, confounding factors and common oncogenes. The use of such gene sets, thus, does not provide the advantage of gaining knowledge and insight into the actual mechanisms of TMM pathway activation. We think that the TMM pathways constructed with this methodology and the PSF based computation of their activities is more valuable in this regard.

It is important to mention that the TMM activity values computed by PSF are strictly dependent on the phenotypic composition of the data, since the PSF algorithm takes as input the fold change values that are computed in reference to the mean expression value of each gene across the samples, as described in Methods.

One should be cautious in interpreting the biological meaning of the TMM pathways generated by this framework. The genes and interactions included in the pathways are selected based on their ability to aid in classification of TMM phenotypes. If a gene participates in both pathways or has extra-TMM roles as well, it will not add to classification power of the pathways. Otherwise, the gene can be of exclusive importance for the pathways, but its involvement in the TMMs may be governed not by its expression but by its post-transcriptional modifications or recruitment to telomeres, which will again disfavor its inclusion in the TMM pathways. This essentially means that the TMM pathways include genes which participate in the cells decision to choose between ALT or Telomerase TMMs and might not fully describe the telomere maintenance mechanisms. Finally, we have limited the networks to the last molecular events leading to TMM activation: one could go deeper and gather information about more upstream interactors, but this requires bigger datasets, as the network becomes bigger and more complicated at upstream levels.

## Conclusions

In conclusion, we have constructed TMM pathways with classification power to distinguish between ALT positive/negative and Telomerase positive/negative phenotypic states based on gene expression data. These pathways allow for further extensions in hand with accumulation of data and knowledge on TMM mechanisms. As new data come in, researchers may extend or modify the existing models to see whether their prediction accuracy increases. Therefore, the presented methodology is intended for two purposes: *in silico* modeling of TMM signaling in order to gain deeper understanding of telomere length maintenance processes and their regulation on one hand, and assessment of TMM phenotypes from gene expression data on the other hand. Even though we have used here only gene expression data to assess the activities of the TMM pathways, other layers of information would add to the accuracy of classification, such as genomic variations, abundance of telomeric repeat variants, as well as DNA methylation and histone modification data.

## Materials and Methods

### Datasets

In order to construct valid TMM pathways, able to predict the TMM phenotype of a cell/tissue from gene expression data, we have obtained samples with experimentally validated TMM phenotypes for which gene expression data was available. For this purpose, we have downloaded the datasets deposited by Lafferty-Whyte *et al* in the Gene Expression Omnibus (GEO) repository [77]. These datasets contain microarray gene expression profiling data for ten cell lines of different tissue origin, and seventeen liposarcoma tumor samples along with four human Mesenchymal Stem Cells (hMSC) isolated from the bone marrow of healthy individuals (GEO accession: GSE14533). Among the cell lines, four were ALT positive (ALT^+^/Telomerase^−^), four were telomerase positive (ALT^−^/Telomerase^+^), and two didn’t have any active TMM (ALT^−^/Telomerase^−^). Among the liposarcoma samples, nine were ALT positive (ALT^+^/Telomerase^−^), as assessed by the presence of APBs, and eight were telomerase positive (ALT^−^/Telomerase^+^), as tested by the telomeric-repeat amplification protocol for telomerase activity detection [78].

### Pathway reconstruction and extension

The pathways were constructed based on protein-protein, protein-RNA interactions that lead to activation of the ALT and Telomerase TMMs, curated from the literature. The pathway maps were built in the Cytoscape environment (http://cytoscape.org/)

Each node in the pathway represents a gene product (protein or RNA), a protein complex, or a process. The edges in the pathway describe functional associations between the nodes. Those are of two types: activation and inhibition, depending on the effect of the source node on the functionality of the target node. It should be noted that the associations in the pathway refer to functional effects of direct protein-protein, protein-RNA interactions between the nodes: indirect effects or regulatory effects on expression are not considered.

We have first constructed initial pathways based on genes with clearly established roles in either TMM, and assessed their ability to predict TMM phenotypes of the studied samples. We then extended the core pathways iteratively, by adding nodes and changing their positions in the pathways. We have evaluated the prediction power of the TMMs at each extension step to arrive at the ‘final’ pathways with best scores (Figure 1). Measures for prediction accuracy are described in detail below.

### Data preprocessing

The gene expression values for technical replicates were averaged. For multiple gene mappings per microarray probe, the probe with highest standard deviation of values was considered. The cell lines and tissue samples (liposarcoma tumors and hMSCs) were further processed in separate sets. For each set, the gene expression values were converted to fold change (FC) values by dividing the expression of each gene to its mean expression across the samples. FC values higher than the 99.5% percentile were set to the value at that percentile. These FC values were then mapped onto pathway nodes and used for assessment of pathway activity (pathway signal flow analysis, see below).

### Pathway signal flow analysis

We have developed a Cytoscape app called TMM for pathway signal flow calculations and for generation of automatic reports on pathway performance (http://apps.cytoscape.org/apps/tmm). For Pathway Signal Flow calculation, TMM app makes calls to the PSFC app for Cytoscape [81] using its default parameters and the signal propagation rules described below. The Pathway Signal Flow (PSF) algorithm [81–83] computes the strength of the signal propagated from the pathway inputs to the outputs through pairwise interactions between nodes, based on their fold change expression values. For each source-target interaction, the FC values are multiplied for edges of type ‘activation’ (FC_source_ * FC_target_) and divided for edges of type ‘inhibition’ (1/FC_source_ * FC_target_). The weighted sum of multiple signals from many sources is assigned as the signal at the target node (the ‘proportional’ option in PSFC). The signal propagation starts from input nodes, spreads through the intermediate nodes and arrives at the sink nodes. In our case, there is a single sink node for each TMM pathway (labeled “ALT” and “Telomerase” respectively). The higher the initial FC value of a node, and the more activation signals it gets from upstream nodes, the higher will be its activity value (or PSF score), and vice versa. The PSF algorithm returns a single PSF score for each node in a pathway. We are, however, interested only in the sink nodes of each TMM pathway: the ‘Telomerase’ and the ‘ALT’ nodes. The PSF scores of these nodes reflect the overall activity of the pathways.

We have upgraded PSFC to also apply functions onto protein complexes: if involvement of all the subunits is obligatory for the complex to function, then the minimum signal of all the subunits is assigned to the complex, otherwise the complex is treated with PSF default rules. Note that nodes where gene expression values were not available were omitted in calculations and assigned a PSF value of 1.

The PSF algorithm is calibrated in a way that if all the nodes in a pathway have FC values of 1, that produces unity PSF values at sink nodes. In our case, the FC value for each gene is computed by taking as a reference the average expression across the samples. This leads to FC values higher or lower than 1, depending on the differential expression of the gene across samples with different TMM phenotypes on one hand, and on the phenotypic composition of the studied samples on the other. Thus, while high PSF values indicate on higher activity of the TMM pathway, there is no predefined PSF threshold that classifies the pathways to be either ‘active’ or ‘inactive’.

### Fitness function

Given gene expression data of a set of samples with either ALT^+^/Telomerase^−^, ALT^−^/Telomerase^+^, ALT^−^/telomerase^−^ or ALT^+^/telomerase^+^ phenotypes, the TMM app is be able to classify those based on the TMM pathway activities, or PSF values. Ideally, ALT^−^/Telomerase^+^ samples should have high PSF values for the Telomerase pathway and low PSF values for the ALT pathway. ALT^+^/Telomerase^−^ samples should score high on the ALT pathway, and low on the Telomerase pathway. The samples with ALT^−^/telomerase^−^ phenotypes should score low at both pathways. This is reflected by the so called fitness function value that consists of two types of metrics.

The first metric is based on median difference of PSF values for ALT and Telomerase pathways between the studied phenotypic groups. The higher the difference, the more is the more the fitness function value. The significance of these mean differences was tested with Kruskal-Wallis rank sum test.

The second metric is based on the accuracy of PSF-based classification of the four possible TMM phenotypic groups. As it was mentioned above, there is no PSF threshold to define if the pathway is ‘active’ or ‘inactive’. Thus, we have used linear support vector machines (SVM), to divide the samples into ALT^+^ or ALT^−^, and Telomerase^+^ or Telomerase^−^ groups for each TMM, respectively. We have then plotted the samples onto a 2D space, where Telomerase PSF values were on the *x* axis, and ALT PSF values were on the *y* axis. The classification accuracy was then assessed based on the ratio of correct versus incorrect classifications.

Computation of these performance metrics and generation of respective plots has been implemented in the TMM app (see below).

For cell lines we possessed three phenotypic groups: ALT^+^/Telomerase^−^, ALT^−^/Telomrase^+^ and ALT^−^/Telomerase^−^. For liposarcoma tumors, we had two phenotypic annotations: ALT+/Telomerase− and ALT−/Telomerase+. We have not included hMSCs, since no evidence exists for hMSCs to have either of the TMM mechanisms activated: according to some reports they have low (or not detectable) levels of telomerase activity [79] and no signs of ALT activity [80]. It should also be noted that even though it’s possible for ALT and telomerase dependent TMMs to coexist in the same cell [39], all the cases with ALT^+^/Telomerase^+^ classifications in our datasets were considered as incorrect. This is reasoned by that fact that none of the samples in our datasets were annotated to possess both of the phenotypes.

### TMM app design and implementation

The TMM software is written in Java (major version 8) and implemented as an app for Cytoscape. It possesses three core functionalities (1) fold change value calculation for TMM genes based on gene expression data supplied by the user, (2) PSF activity calculation of the TMM pathways based on command calls to the PSFC app (version 1.1.4 or higher) for Cytoscape, and (3) generation of statistical reports and plots. Visualization of pathways and their PSF flows is also provided by the app in Cytoscape environment.

The user may either have experimental annotations of assumptions on TMM states, or they may want to use the app for assessment of otherwise unknown TMM phenotypes of their samples. We will refer to these cases as *Use case A* and *Use case B* respectively.

### Use case

#### Supply TMM network

The user should first load the TMM network (containing TMM pathways) in the Cytoscape environment. Pathways may be downloaded from the TMM project website (http://big.sci.am/software/tmm/), and can further be modified by the user. Modifications of pathways may be made by adding novel genes, which the user thinks might be of functional importance, by deleting the existing genes, or by changing the interactions and the place of the gene in the pathway topology. Before performing any modifications to the pathways the user is advised to visit the http://big.sci.am/software/tmm/tmm-genes/ page and see if they find any information about the gene of interest: whether it’s already been tested or not yet. The user may also want to refer to the citations provided for each interaction in the Network Edge Table in Cytoscape.

#### Supply the gene expression data

The gene expression data is supplied to the TMM app, which then computes the fold change values relative to the gene-wise global means across samples. The user may first want to preprocess the gene expression data according to the recommendations available in the TMM user guide (http://big.sci.am/software/tmm/#userguide).

#### Use case A: supply TMM labels

The user may indicate the experimentally tested or assumed TMM phenotypes of the samples, which will allow TMM to generate statistics reports and color-labeled plots.

#### Use case B

If no TMM phenotypes are available or assumed, the user will get TMM activity distributions across samples.

#### Results generation and visualization

After supplying the data and labels (for *Use case A*) the user clicks the button to run the pipeline. TMM app makes a call to the PSFC app through Cytoscape’s command interface. PSFC generates a summary file with PSF values, which is then read by the TMM app. The latter generates graphical and text reports based on this summary file. Besides the report files, the user may also observe the coloring of the nodes in the pathways based on their PSF values in Cytoscape.

Example archives containing network and input files and the reports are available at http://big.sci.am/software/tmm/#downloads.

